# The Contribution of Baseline Circulating Endocannabinoids to Individual Differences in Human Pain Sensitivity: A Quantitative Sensory Testing Study

**DOI:** 10.1101/2025.08.22.671762

**Authors:** S. A. Fatemi, S. S. Abssy, S. L. Bourke, K. B. Murray, C. Kyeremaa-Adjei, L. Honigman, N. Mohabir, C. Sexton, M. A. Cormie, R. Tomin, I. Boileau, L.Y. Atlas, D.P. Finn, M. Moayedi

## Abstract

The endocannabinoid (eCB) system—comprising cannabinoid receptors, eCBs (anandamide— AEA, 2-arachidonoylglycerol—2-AG) and related *N*-acylethanolamines (NAEs; *N-*palmitoylethanolamide—PEA, and *N-*oleoylethanolamide—OEA), and metabolizing enzymes (e.g., fatty acid amide hydrolase; FAAH)—modulates nociceptive circuits in rodents. In humans, the *FAAH* C385A polymorphism is associated with reduced pain sensitivity, suggesting eCB tone influences individual pain differences, but this has yet to be tested. Here, we determined whether the eCB system is associated with somatosensory and pain sensitivity measured with quantitative sensory testing (QST) in 91 healthy participants (39 males, 52 females). We tested three hypotheses: (1) *FAAH* C385A polymorphism, cannabis use, and sex affect serum eCB/NAE concentrations; (2) *FAAH* C385A carriers show altered pain sensitivity versus non-carriers; and (3) baseline serum eCB/NAE concentrations are associated with QST measures. eCB/NAE concentrations were not statistically different based on sex (*p* > .05), based on *FAAH* genotype (*p* > .05), and based on cannabis use (*p* > .05). To address collinearity of AEA, OEA and PEA, we performed a principal components analysis, which identified a single component of FAAH substrates. Linear regressions found that *FAAH* genotype did not affect QST measures and that baseline 2-AG and FAAH substrate concentrations were not associated with QST measures, except pressure pain thresholds (PPT; *p* = 0.003), which were associated with AEA and OEA. Baseline eCB/NAE levels and FAAH genotype are not associated with the outcome measures of standard QST tests that rely on point estimates in healthy adult humans; nonetheless, circulating FAAH substrate levels were associated with PPT.

## INTRODUCTION

There are significant individual differences in human pain sensitivity [20,23,37,76]. Biological factors such as sex, the state of the nociceptive and pain modulatory systems, and genetic variability contribute to individual differences in pain [6,21,23,24,30,59]. Neuromodulators that affect nociceptive pathways and descending modulatory circuits may thus contribute to individual differences in pain sensitivity. One such neuromodulatory system identified in preclinical models of pain is the endocannabinoid (eCB) system [25,32,39,46,94,103].

The eCB system includes cannabinoid receptors; endogenous cannabinoid ligands, including 2-arachidonoylglycerol (2-AG), *N-*arachidonoylethanolamine/anandamide (AEA), and related *N*-acylethanolamines (NAEs) including *N-*palmitoylethanolamide (PEA) and *N-*oleoylethanolamide (OEA); and metabolizing enzymes, including fatty acid amide hydrolase (FAAH) and monoacylglycerol lipase. Although NAEs are not strictly eCBs, they may also have pain modulatory effects, both directly through peroxisome proliferator-activated receptors [85], and indirectly via elevation of AEA levels through substrate competition at FAAH [4,52,60,78].

Preclinical research has shown that eCBs and NAEs are present at every level of the nociceptive system, including within key structures of descending modulatory circuits [13,39,44,79,80,94,103]. In rodents, eCBs modulate rodent thermal and mechanical sensitivity [27,66,67]; for a comprehensive review, see [32]. For example, FAAH inhibition reduces carrageenan- and lipopolysaccharide-induced mechanical and thermal sensitization [54,65,74]. Further, there are sex differences in eCB ligand concentrations in the nociceptive and modulatory pathways [62] and effects on pain [10,99]. However, the contribution of eCBs to pain sensitivity in healthy humans remains unknown.

Evidence supporting the role of the eCB system in human pain comes from studies of the functional *FAAH* rs324420 (C385A) single nucleotide polymorphism (SNP), which can result in elevated AEA, OEA and PEA levels [69,92]. The global frequency of the CC, AC and AA genotypes are 55.5%, 36.7% and 7.8%, respectively (Table S1) [33]. Variants of the *FAAH* gene are associated with lower cold pain sensitivity, reduced postoperative analgesic needs compared to non-carriers [17], and congenital pain insensitivity [107]. Further, CB^1^ receptor antagonism selectively reduced non-opioidergic placebo analgesia [9,83]. Thus, natural eCB system variation may account for individual pain sensitivity differences. However, circulating eCB concentrations are affected by cannabis use [51], and may thus modify the relationship with pain sensitivity, but this remains unknown. Individual differences in pain sensitivity can be measured with quantitative sensory testing (QST) [55,87-89]—a standardized set of psychophysical tests to assess multiple sensory modalities (innocuous and noxious thermal and mechanical stimuli).

Here, we explore whether the eCB system is associated with individual QST measures. We first explore whether the *FAAH* C385A SNP, sex, and cannabis use affect circulating eCB/NAE levels. Based on the literature, we expect that those with the C385A SNP have higher levels of FAAH substrates (AEA, OEA, and PEA) [22,91,93], and that females have lower levels of FAAH substrates than males [3,62]. Next, we test whether individuals with the *FAAH* C385A polymorphism show different pain sensitivity than individuals without the polymorphism. We hypothesize that A-allele carriers have lower pain sensitivity. Third, we test relationships between baseline circulating eCB/NAE levels and QST measures, hypothesizing that higher eCB/NAE concentrations correlate with lower pain sensitivity. Finally, we explore whether there are sex differences in these relationships, anticipating that there will be differences, but without directional predictions.

## METHODS

### Participants

One hundred and ten healthy adults (62 females and 48 males) aged between 21 and 45 years of age were recruited from the University of Toronto and surrounding community between October 6, 2022, and June 30, 2024. All participants consented to procedures approved by the University of Toronto’s Human Research Ethics Board (Protocol #38322). The sample size we aimed to recruit was 100 participants, as this was the first study of its kind.

Participants were screened against the following exclusion criteria with self-report: current pain, history of chronic pain, chronic illness, psychiatric disorder or neurological disorders, pregnancy, breastfeeding, immunocompromised, specific dietary restrictions (that would not allow participants to eat the carbohydrate snack provided, see below), < 20 years of age, and ≥ 45 years of age (to mitigate any effects of cutaneous sensory changes that occur with aging, and can affect pain sensitivity).

Participants were asked to refrain from any cannabis-use 24 hours prior to the session, to minimize the effects of Δ^9^-tetrahydrocannabinol (Δ^9^-THC) on pain. Given the high prevalence of recreational cannabis use in Canada following legalization [48], complete exclusion of cannabis users was deemed impractical. Additionally, as dietary fats lead to the synthesis of some eCBs and fatty acid amides, participants were asked to fast overnight (for 12 hours) prior to the study session. To control for the dynamic levels of circulating eCBs and NAEs that exhibit a circadian rhythm [49], all the sessions took place between 9:00AM and 12:00PM.

### Experimental Session

The data presented in this study are part of a larger study with an MRI scanning session. The MRI data are outside the scope of this study. We describe the entire experimental session for completeness.

Upon arrival to the Centre for Multimodal Sensorimotor and Pain Research, University of Toronto, and after consent, fasted participants were given the choice of a 30g blueberry oat bar or 30g chocolate chip oat bar (MadeGood© Mornings, Riverside Natural Foods, Toronto, ON, Canada) to control for the nutritional value of the snack. Participants then completed two additional screening questionnaires: the Beck Depression Inventory II (BDI-II) and the Mini Mental State Exam (MMSE). Participants then completed a battery of questionnaires (see *Questionnaires* below), provided a blood sample, and finally underwent a QST battery. Figure 1 provides a summary of the experimental session. A subset of 99 participants was then taken to the MRI facility by taxi and underwent a brain imaging session (outside the scope of this study). Participants were financially compensated for their participation in the study.

**Figure 1:**
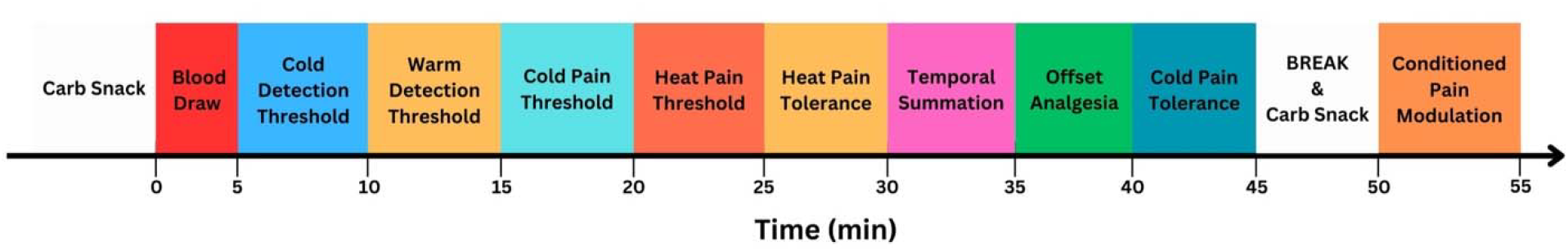
Schematic representation of the experimental session. Note that the offset analgesia and conditioned pain modulation are not included herein as they are the focus of another study.

### Screening Questionnaires

The BDI-II is a 21-item self-report questionnaire that allows individuals to rate their agreement with statements about thoughts and feelings in daily life and is used to measure attitudes and symptoms of depression [8]. Participants who scored > 20 on the questionnaire, demonstrating moderate depression were briefly consulted to ensure that they are feeling well enough to proceed with the experiment, and if they agreed to continue, they were included in the study. Those who indicated suicidal ideation (score > 1 on item 9) were excluded. We also collected the Patient Health Questionnaire (PHQ-9), a multipurpose instrument that allows screening of depression and serves as a tool to diagnose and monitor the severity of depression [58]. This test supplemented the BDI to screen participants for depression, and to align protocols with other studies to increase the utility of the data when published for open access use.

The MMSE is a tool for systematically and thoroughly assessing mental status. The MMSE is an 11-item questionnaire that can be used to ensure there are no cognitive impairments by measuring five areas of cognitive function: registration, orientation, recall, attention and calculation, and language [40]. This questionnaire served to ensure that participants were alert and could follow directives prior to beginning the experiment.

### Questionnaires and Measures

We collected several questionnaires to capture factors associated with pain sensitivity or circulating eCB/NAE concentrations. Related to the current study, participants completed a demographics questionnaire, and the self-report Daily Sessions, Frequency, Age of Onset, and Quantity of Cannabis Use Inventory (DFAQ-CU) [29]. The demographics questionnaire captured participants’ age, sex, gender, and highest level of education attained. The DFAQ-CU is a psychometrically sound inventory which assesses the frequency, age of onset, and quantity of cannabis use. In this study the DFAQ-CU was used to assess participants’ history of cannabis use. As the scale does not provide a continuous outcome measure, we summarized each participant’s score on an ordinal scale: 0 = never used cannabis; 1 = previously used cannabis, but not in the last 3 months; 2 = currently using cannabis. Participants also had their height and weight measured to calculate body mass index (BMI).

Participants also completed several other questionnaires outside the scope of this study, which we list here for completeness: the Pain Catastrophizing Scale, Fear of Pain Questionnaire– III, Positive and Negative Affect Scale, Action Control Scale, the PROMIS Anxiety Short Form 7a, and the State-Trait Anxiety Inventory.

### Blood Sample Collection and Processing

Blood samples were collected from the participants by certified phlebotomy technicians. Two vials of blood were collected: 2 mL sample into BD Vacutainer® spray coated K2EDTA 3.6 mg blood tubes, and a 6 mL sample in a BD Vacutainer® Serum tube. The samples were gently inverted 10 times following collection. The 2 mL sample was immediately snap frozen on dry ice and kept at - 80°C until shipment. The 6mL sample was left to clot at room temperature for 30 minutes. Next, this sample was centrifuged at 1400 g for 15 minutes at 4°C. The serum layer was collected with a disposable transfer pipette and aliquoted into 2 labelled 1.5 mL microcentrifuge tubes. The samples were snap-frozen on dry ice and kept at -80°C until analysis. Samples were shipped in dry ice to the University of Galway, Ireland for eCB/NAE quantification and FAAH C385A genotyping.

#### Circulating Endocannabinoid Quantification

Serum eCBs and NAEs were quantified using liquid chromatography-tandem mass spectrometry (LC-MS/MS) as described previously, with some modifications [7,14]. Briefly, 20 μL 100% acetonitrile containing deuterated internal standards (50 ng of 2-AG-d8, 2.5 ng of AEA-d8, 2.5 ng of OEA-d4 and 2.5 ng of PEA-d4) were added to each sample. Samples were then vortexed and allowed to equilibrate for 10 minutes on ice. For protein precipitation, 1 ml of 100% ACN containing 0.1% formic acid, maintained at 4°C, was added to each sample and immediately vortexed and incubated on ice for 30 minutes. The precipitated proteins were pelleted by centrifugation at 18,620 g for 15 minutes at 4°C. A filter (FisherbrandTM Non-sterile PTFE Hydrophyl, 25 mm, 0.45 µM Syringe Filter, Fisher Scientific, Ireland) was placed in a 5 ml SafeSeal tube (SARSTEDT, Ireland) for each sample. 1 ml syringes were attached to the filters by carefully inserting the syringe neck into the filter inlet. The plunger/piston from each 1 mL syringe was removed and saved in a sterile container. 1100 µL of the supernatant was removed without disturbing the pellet and loaded into the syringe. The supernatant was allowed to flow through the filter until the syringe was emptied. Once the supernatant had flown through the syringe into the microcentrifuge tube, 700 µL of 100% ACN was added to the syringe to displace the dead volume of the filter. The syringe plunger was slowly re-inserted, and the 100% ACN was pushed through the filter. The collected eluate was vortexed, and 500 µL of the total volume was transferred to a new 1.5 mL microcentrifuge tube and dried down at 45°C for approximately 1 hour in a centrifugal concentrator (Eppendorf Concentrator plus complete system, Davidson and Hardy Ltd, Ireland). Each sample was reconstituted in 20 µL of 100% ACN before transferring it to HPLC vials and were then separated on a Zorbax® C18 column (150 × 0.5 mm internal diameter; Agilent Technologies, Cork, Ireland) by reversed-phase gradient elution initially with a mobile phase of 65% acetonitrile and 0.1% formic acid, which was ramped linearly up to 100% acetonitrile and 0.1% formic acid over 10 min. Under these conditions, 2-AG, AEA, PEA and OEA eluted at the following retention times: 6.6 min, 6.9 min, 7.0 min, and 7.2 min, respectively. Analyte detection was carried out in electrospray-positive ionization mode on an Agilent 1260 Infinity II HPLC system coupled to a SCIEX QTRAP 4500 mass spectrometer operated in triple quadrupole mode (SCIEX Ltd, Phoenix House Lakeside Drive Centre Park, United Kingdom). was performed by ratiometric analysis and expressed as nmol or pmol per mL of serum.

#### FAAH C385A Genotyping

Genomic DNA (gDNA) were extracted using standard protocols (Nucleospin® DNA isolation kit, Macherey-Nagel, Düren, Germany). All samples were genotyped for the C385A *FAAH* SNP (rs324420) using predesigned TaqMan primers and universal genotyping master mix (Life Technologies). Genotyping assays were performed according to manufacturers’ protocols using an Applied Biosystems (ABI) StepOne Plus PCR machine and ABI allelic discrimination software.

### Quantitative Sensory Testing

#### Thermal QST measures

##### Innocuous Thermal Detection and Pain Thresholds

We measured the participant’s ability to detect thermal stimuli in the innocuous and noxious ranges. We used the T11 thermode with a stimulation area of 9 cm^2^ to generate thermal sensations in the cool, warm, noxious cold and noxious heat ranges on the volar forearm with the Thermal Cutaneous Stimulator device (TCS II, QSTLab, Strasbourg, France) [41,105,106]. The thermode was placed on the right volar forearm and delivered thermal stimuli on the participant’s skin to evaluate cool detection threshold (CDT), warm detection threshold (WDT), cold pain threshold (CPT) and heat pain threshold (HPT). For each modality, the thermode started at a baseline temperature of 30°C and increased or decreased at a rate of 1°C/s for CDT, WDT, CPT and HPT protocols [34]. Participants were asked to press a response button when a change in the perception towards the target stimulus was detected for the first time. For innocuous stimuli, we obtained four measurements for each detection threshold, with an interstimulus interval of 2 seconds. The first trial was discarded in all participants. For pain detection thresholds, we provided participants with the International Association for the Study of Pain’s definition of pain: “an unpleasant sensory and emotional experience associated with, or resembling that associated with, actual or potential tissue damage” [86]. We obtained three measurements for each thermal pain threshold, and all three were included in the analysis. The arithmetic mean was calculated for each thermal detection threshold (in °C). For safety and technical reasons, cold pain thresholds were considered maximal at 0 °C, and heat pain thresholds were considered maximal at 50 °C.

##### Heat Pain Tolerance (HPTol)

Participants underwent a HPTol paradigm with the thermode placed on the right volar forearm of the participant. The stimulus increased at a rate of 1°C/s. Participants were instructed to press the response button whenever they could no longer tolerate the heat pain elicited by the thermal probe. The TCS device was programmed to have an upper limit of 50°C for safety. Note, that as HPTol was added to the battery of tests after the study had started, n = 11 did not undergo this test and the sample size for the HPTol test was n = 88.

##### Cold Pain Tolerance (CPTol)

CPTol was measured with the cold pressor task. An 18 L water-bath (FBATH18, Techne, Burlington, NJ, USA) with a digital immersion circulator (FTE10DPC, Techne) and a low-temperature cold-pressor with a rigid coil probe (FRU2P, Techne). The water bath was set to 8 ± 1°C. Participants immersed their dominant hand into the bath up until their wrist with their palm facing down and fingers spread out. A timer was set at time of immersion. Participants were instructed to keep their hand in the water as long as possible, up to the point they could no longer tolerate the cold pain. They were instructed to verbally announce “pain” when the sensation of the cold water became painful, and this time was recorded as a cold pain threshold. Once participants removed their hand, they provided a pain intensity rating on a verbal numeric rating scale, rating from 0 (no pain) to 100 (worst pain imaginable). The duration of immersion was set to a maximum time of 180 seconds if participants could fully tolerate the stimulus. CPTol was calculated by subtracting the time of pain threshold from the time at which pain tolerance was maximized.

#### Mechanical QST measures

##### Pressure Pain Threshold

PPT was assessed using a handheld digital algometer (Algomed) fitted with a 1 cm^2^ probe. With participants seated upright, the probe was applied manually to three adjacent sites on the dominant upper trapezius. Participants held a response button in their non-dominant hand and were instructed to press it and say “pain” as soon as the sensation changed from pressure to pain. Each of the three trials was separated by approximately 10 seconds, and the PPT value was defined as the arithmetic mean of these three measurements.

##### Temporal Summation of Pain (TSP)

TSP was conducted on the dominant volar forearm. Three distinct locations no further than 2 cm apart were marked in a triangular shape. Participants were instructed to close their eyes. A single weighted pinprick stimulator of 256 mN with a 0.25 mm diameter contact area (MRC Systems GmbH-Medizintechnische Systeme, Heidelberg, Germany) was used to perform a single pinprick stimulus. Participants reported their perceived pain intensity on a verbal numerical rating scale between 0 (no pain) to 100 (worst pain imaginable). Next, a train of ten 256 mN pinprick stimulations was applied at 1 Hz to the two other marked locations. Following the training of ten repeated stimuli, participants rated their perceived pain from the last stimulus. A 2-minute break was then taken prior to stimulation of the next site. This procedure was repeated a total of three times. The final value for TSP was calculated by taking the difference between each of the ratings after the tenth stimulus and the ratings after the first stimulus and then computed an arithmetic average of those differences.

### Statistical Analyses

All statistical analyses were performed in JASP v.0.19.3[96], unless otherwise noted.

#### Differences in eCB/NAE ligands based on sex, FAAH C385A SNP genotype, and cannabis-use

To determine whether there are sex differences in levels of circulating eCBs (AEA, 2-AG) and NAEs (OEA, PEA) we first tested the residuals for normality with Shapiro-Wilk’s test, with significance set at *p* < 0.05. Normally distributed variables were compared between sexes with an independent samples *t*-test. Non-normally distributed variables were compared with a Mann-Whitney *U* test. Significance was set at *p* < 0.05.

Similarly, to test whether the *FAAH* C385A SNP had a significant effect on levels of circulating eCBs/NAEs, all eCB/NAE concentrations were first standardized (z□scored) across participants. We then tested the residuals for normality with Shapiro□Wilk’s test, with significance set at *p* < 0.05. Normally distributed variables were compared between those without vs. those with an A allele with an independent samples *t*-test. Non-normally distributed variables were compared with a Mann-Whitney *U* test. Significance was set at *p* < 0.05.

Next, we sought to determine whether cannabis-use was associated with baseline levels of circulating eCBs/NAEs. We performed a multivariate analysis of variance (MANOVA) with Pillai’s test, to determine whether z-scored AEA, OEA, PEA and 2-AG were different between those who have never used cannabis, those who have previously used cannabis, but do not currently use, and those who currently use cannabis, based on the DFAQ-CU. Significance was set to *p* < 0.05.

#### Data Reduction

As all three FAAH substrates were highly correlated, and to reduce the number of multiple comparisons, we performed a principal components analysis (PCA). First, the variables were z-transformed. We ensured that criteria for PCA were met with a Kaiser-Meyer-Olkin Test for Sampling Adequacy (> 0.5) and Bartlett’s test of sphericity (*p* < 0.05). PCs with an eigenvariate > 1 were accepted as a solution. If more than 1 component was identified, these were orthogonalized with a varimax rotation.

#### Linear Regressions

All dependent variables (QST test outcome measures) were z-transformed. To determine whether pain sensitivity is associated with levels of eCBs/NAEs, we performed two sets of linear regressions: (1) thermal stimuli (WDT, CDT, HPT, CPT, HPTol, CPTol) and (2) mechanical stimuli (PPT, TSP). Each linear regression had a different dependent variable (the QST measure), and the same set of predictors. Predictors were tested across five additive models. The first (null) model included age and BMI. BMI was included as there is some evidence suggesting associations with eCBs and NAEs [68]. In the second model, sex was added. The third model incorporated the *FAAH* SNP genotype in the model. Note that as the AA genotype only had 5 participants, we coded the genotype as a binary variable: without (CC) and with (AA and AC) the SNP. The fourth model added the circulating FAAH substrate level composite score and circulating 2-AG levels. The fifth model added cannabis use to the model. This design allows us to test all three hypotheses in a single analysis. Model selection was based on whether the model fit was significant and based on the lowest Aikaike Information Criterion (AIC). All models were checked for collinearity using the variance inflation factor, ensuring it is below a conservative threshold of 2.5. Significance was set to *p* < 0.05, Bonferroni corrected for all other linear regressions in the test set (i.e., *p* < 0.0083 for thermal stimuli; *p* < 0.025 for mechanical stimuli).

In cases were the FAAH substrate composite score was associated with a QST measure, post-hoc partial correlation analyses were performed to determine which FAAH substrates are associated with PPTs, while controlling for age, BMI, sex, FAAH genotype, and 2-AG levels, to best mimic the linear regression analysis. Significance was set at *p* < 0.0083.

## RESULTS

### Participants

We recruited 110 participants for this study. Of these, six were excluded due to inability to draw blood, three due to incidental findings in MRI, and two withdrew from the study, resulting in a cohort of 99 participants (44 males and 55 females). Of these, serum eCB/NAEs data were missing for 5 participants (3 males and 2 females), *FAAH* SNP genotype was not captured in 3 participants (2 males and 1 female) due to technical issues, and thermal detection thresholds were not recorded for 2 males. Furthermore, as the HPTol test was added after the start of the study, these data are missing for 11 participants (7 males and 4 females). Our final sample included 81 participants for HPTol, 89 participants for WDT and CDT, and 91 participants for additional analyses (see Figure 2). See Table 1 for Demographics information.

**Table 1.**
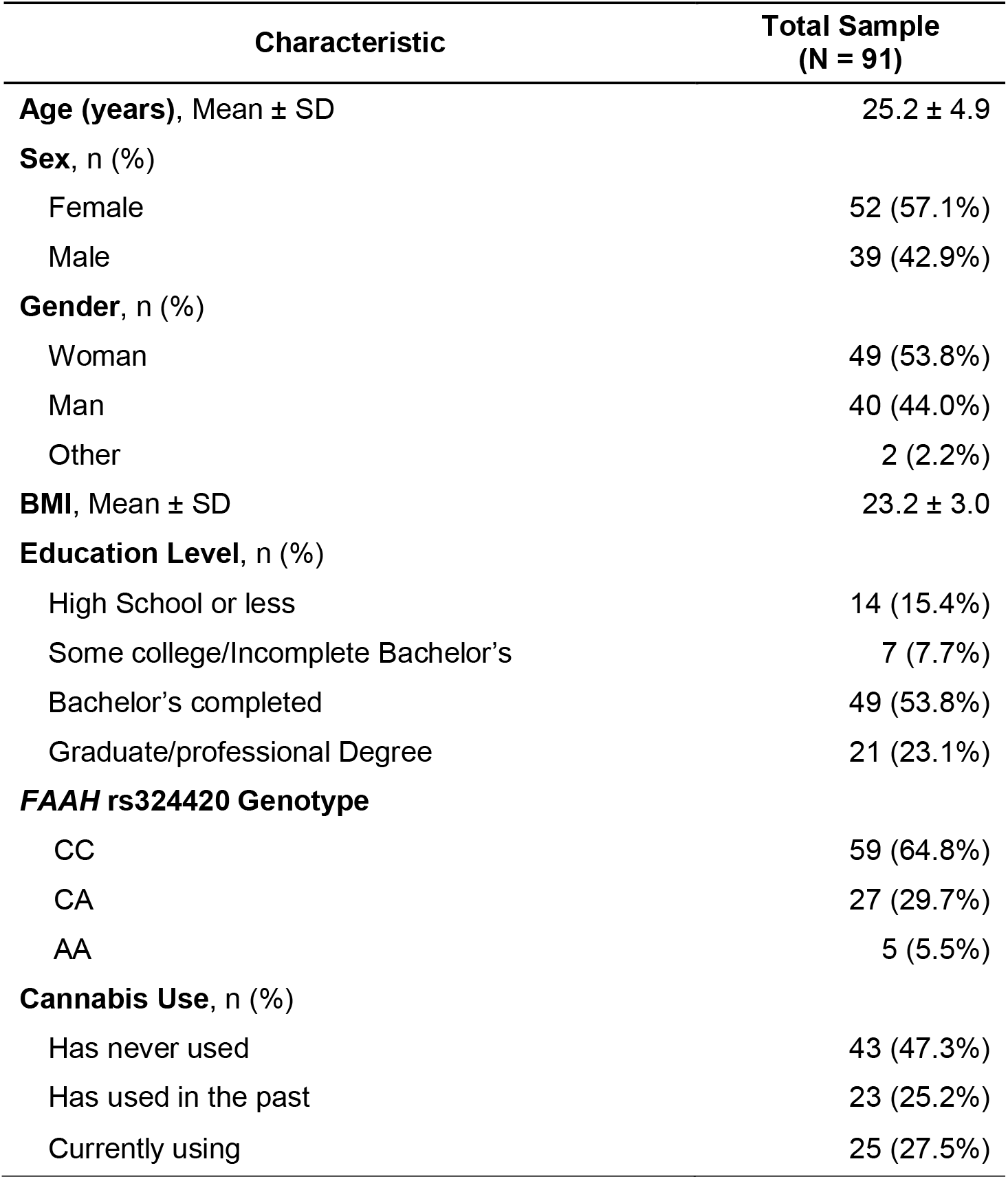
Demographic and anthropometric characteristics of the study sample.

| Characteristic | Total Sample<br>(N = 91) |
| --- | --- |
| **Age (years)**, Mean ± SD | 25.2 ± 4.9 |
| **Sex**, n (%) |  |
| Female | 52 (57.1%) |
| Male | 39 (42.9%) |
| **Gender**, n (%) |  |
| Woman | 49 (53.8%) |
| Man | 40 (44.0%) |
| Other | 2 (2.2%) |
| **BMI**, Mean ± SD | 23.2 ± 3.0 |
| **Education Level**, n (%) |  |
| High School or less | 14 (15.4%) |
| Some college/Incomplete Bachelor's | 7 (7.7%) |
| Bachelor's completed | 49 (53.8%) |
| Graduate/professional Degree | 21 (23.1%) |
| **FAAH rs324420 Genotype** |  |
| CC | 59 (64.8%) |
| CA | 27 (29.7%) |
| AA | 5 (5.5%) |
| **Cannabis Use**, n (%) |  |
| Has never used | 43 (47.3%) |
| Has used in the past | 23 (25.2%) |
| Currently using | 25 (27.5%) |

**Figure 2.**
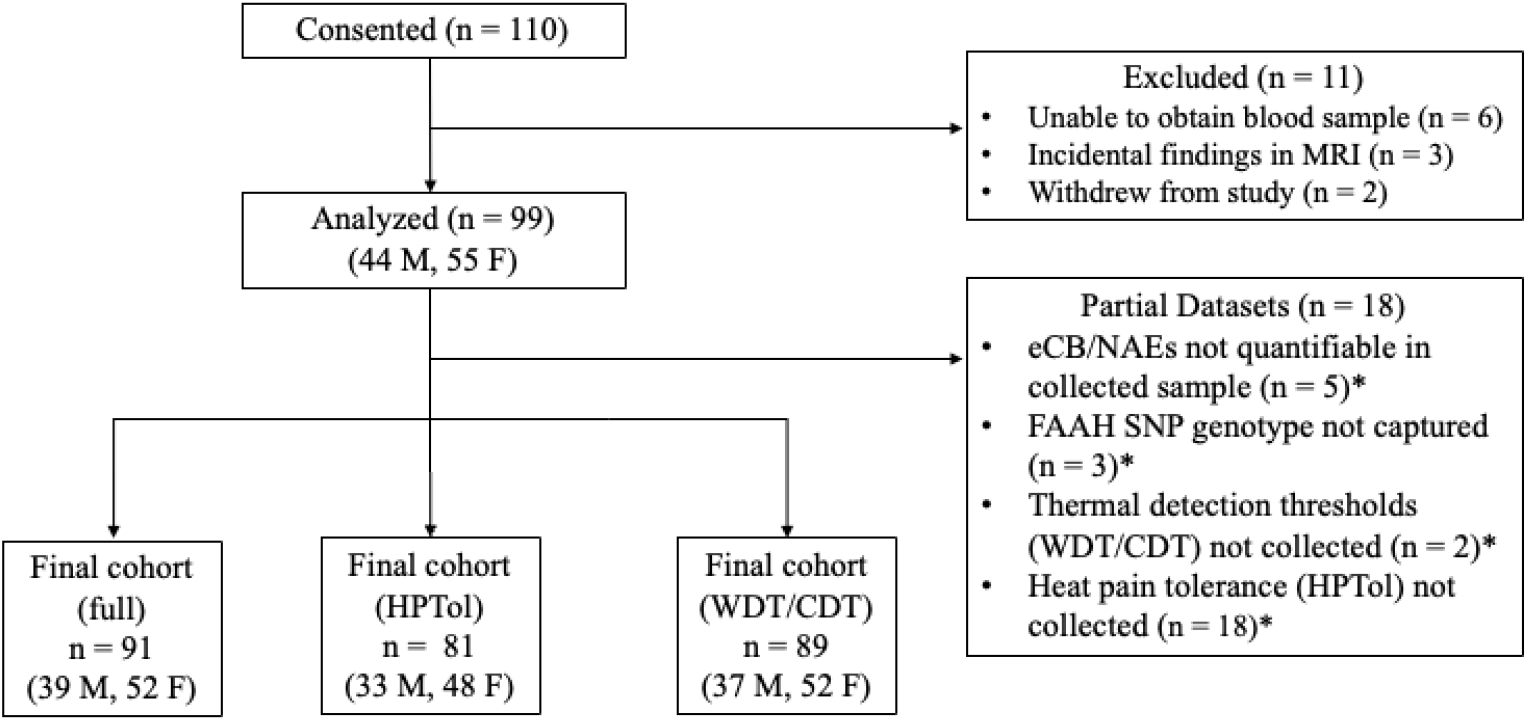
Flow diagram of recruitment and sample. Eighteen participants had partial datasets. *indicates sample overlap.

### No Effect of Sex, *FAAH* SNP Genotype, and Cannabis-use on Baseline Circulating eCBs/NAEs

We found no significant sex differences in baseline levels of circulating eCBs/NAEs (all *p* > 0.12); Table S2). Similarly, there were no differences in baseline levels of circulating eCBs/NAEs based on *FAAH* genotype (all *p* > 0.06; Table S3). Lastly, a MANOVA showed no significant differences in baseline levels of circulating eCBs/NAEs with cannabis-use comparing non-users, previous users, and current users (F^8,172^ = 1.60, Pillai’s trace = 0.14, *p* = 0.13).

### Association between QST measures and eCBs/NAEs

#### Data Reduction

There was a single resultant component (weightings provided in Table S4), which we refer to as “FAAH substrates” herein.

#### Thermal QST Measures

Distributions of thermal QST measures are presented in Figure 3.

**Figure 3.**
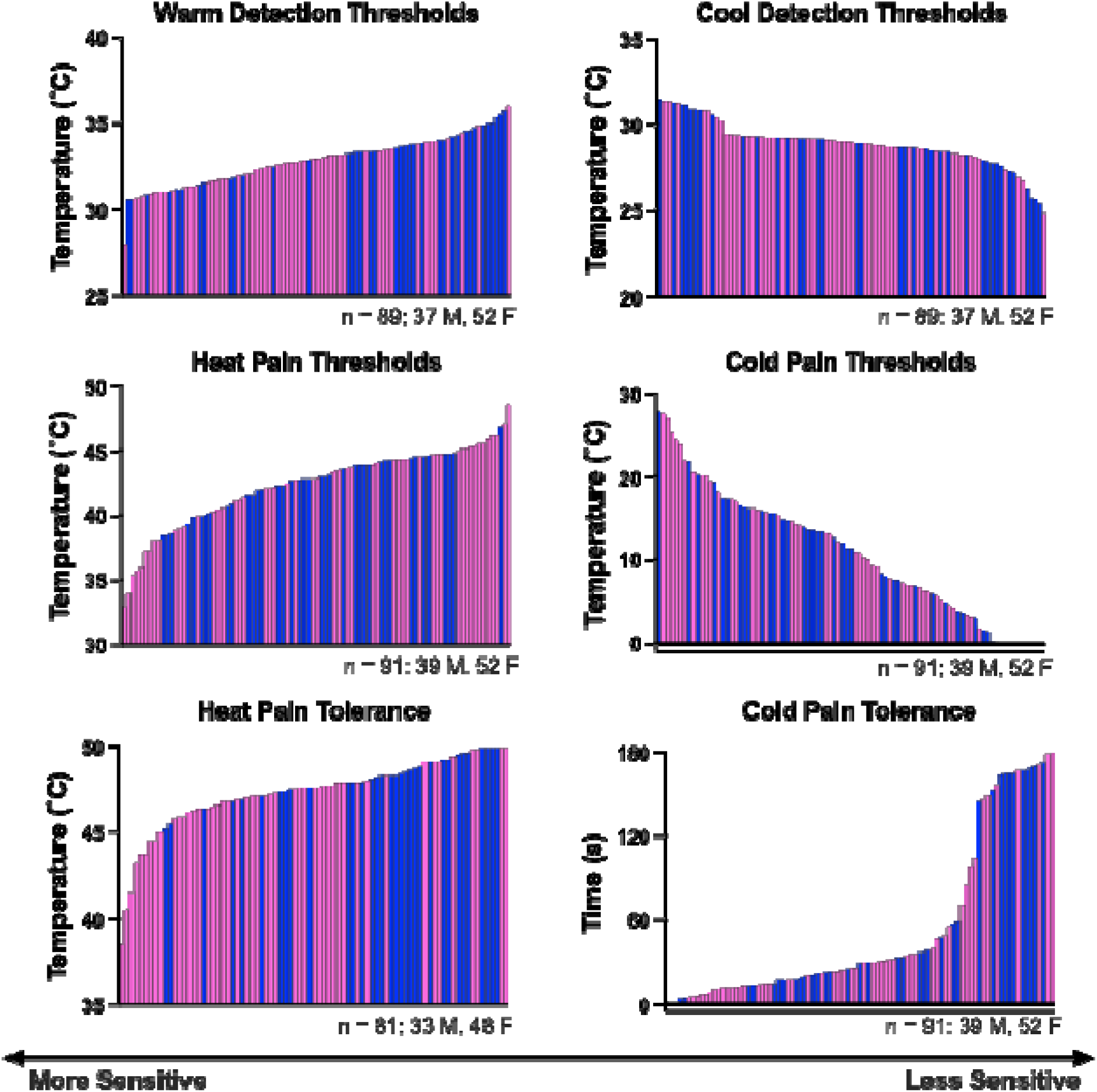
Distribution of Thermal QST measures across participants. Each bar represents a single participant, and participants are ranked in ascending order. Pink bars represent females and blue bars represent males.

##### WDT

Models for WDT are provided in Table S5a. The winning model for WDT was model 1 which included age, sex, and BMI (AIC = 243.0, R^2^ = 0.128, *F*^3,85^ = 4.17, *p* = 0.008). Sex was the only significant predictor in the model (*t* = -2.83, *p* = 0.006, Table S5b), with female sex having lower WDT than males, indicating females were more sensitive to thermal sensations. *FAAH* SNP genotype, circulating FAAH substrates, and cannabis use were not associated with WDT (all *p* > 0.05).

##### CDT

Models for CDT are provided in Table S6. There were no significant models for CDT, indicating that CDT is not associated with sex, *FAAH* SNP genotype, circulating levels of FAAH substrates or 2-AG, or cannabis-use (all *p* > 0.05).

##### HPT

Models for HPT are provided in Table S7. There were no significant models for HPT, indicating that HPT is not associated with sex, *FAAH* SNP genotype, circulating levels of FAAH substrates or 2-AG, and cannabis-use (all *p* > 0.05).

##### CPT

Models for CPT are provided in Table S8. There were no significant models for CPT, indicating that CPT is not associated with sex, *FAAH* SNP genotype, circulating levels of FAAH substrates or 2-AG, and cannabis-use (all *p* > 0.05).

##### HPTol

Models for HPTol are provided in Table S9a. Model 2, which included age, sex, and BMI and *FAAH* allele (AIC = 227.48, R^2^ = 0.176, *F*^*4,76*^ = 4.05, *p* = 0.005) was the winning model. Sex was the only significant predictor in the model (*t* = -2.87, *p* = 0.005, Table S9b), with female sex having lower HPTol than males. *FAAH* SNP genotype, circulating FAAH substrates, and cannabis use were not associated with HPT (all *p* > 0.05).

##### CPTol

Models for CPTol are provided in Table S10. There were no significant models for CPTol, indicating that this measure is not associated with sex, *FAAH* SNP genotype, circulating levels of FAAH substrates or 2-AG, and cannabis-use (all *p* > 0.05).

#### Mechanical QST measures

Distributions of mechanical QST measures are presented in Figure 4.

**Figure 4.**
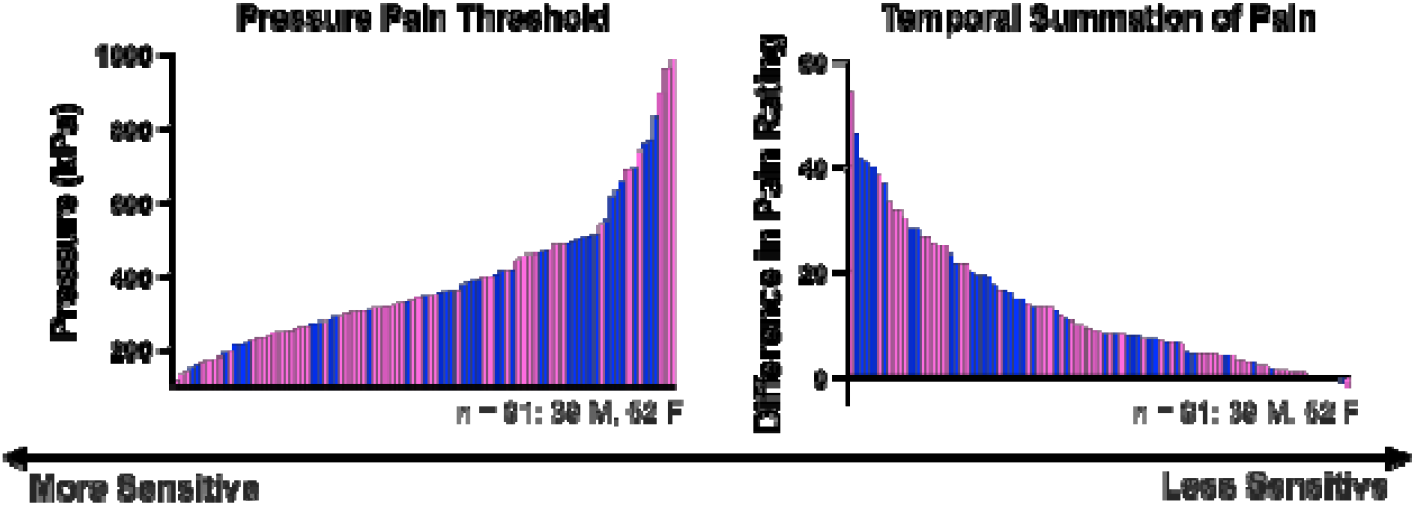
Distribution of mechanical QST measures. Each bar represents a single participant, and participants are ranked in ascending order. Pink bars represent females and blue bars represent males.

**Figure 5.**
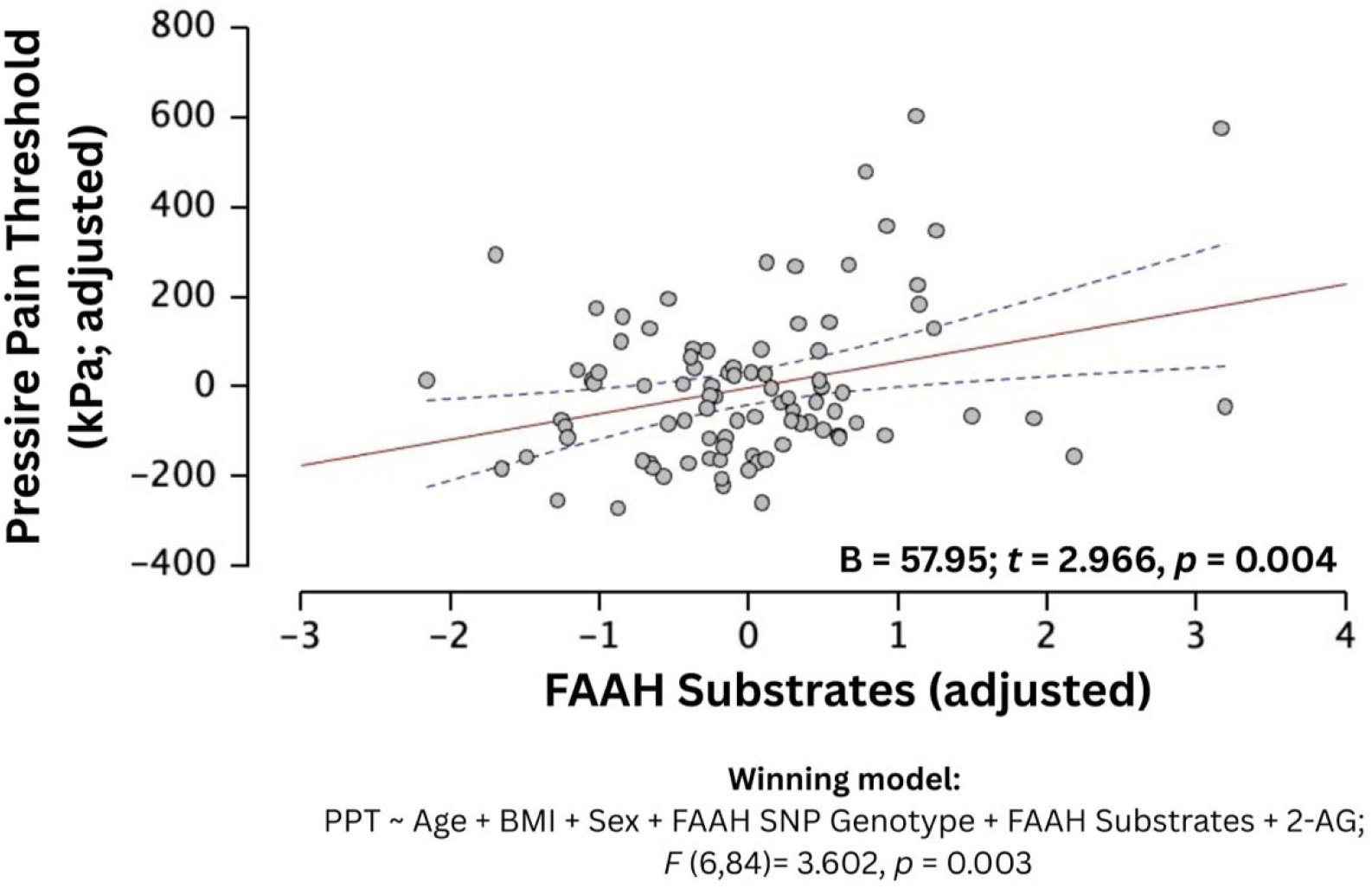
Individual differences in pressure pain threshold are associated with circulating FAAH substrates. Values are adjusted for age, sex, body mass index, 2-AG and FAAH genotype. The regression line is shown in red, and 95% confidence intervals are indicated with dashed blue lines.

##### PPT

Models for PPT are provided in Table S11a. The winning model for PPT was Model 3, which included age, sex, BMI, *FAAH* SNP genotype, circulating FAAH substrates, and 2-AG (AIC = 1202.65, R^2^ = 0.205, *F*^6,84^ = 3.60, *p* = 0.003). The circulating FAAH substrates variable was the only significant predictor in the model (*t* = 2.97, *p* = 0.004, Table S11b), indicating that those with higher levels of FAAH substrates had higher pain thresholds (i.e., less sensitive). *FAAH* SNP genotype, sex and cannabis use were not associated with PPT (all *p* > 0.05).

Post-hoc partial correlation analyses controlling for age, sex, BMI, FAAH genotype and 2-AG revealed that AEA and OEA were associated with PPT (AEA: *r* = .345, *p* = .00099; OEA: *r* = .300, *p* = 0.00494), but PEA was not associated (*r* = .133, *p* = .2234).

##### TSP

Models for TSP are provided in Table S12. There were no significant models for TSP, indicating that TSP is not associated with sex, *FAAH* SNP genotype, circulating levels of FAAH substrates or 2-AG, and cannabis-use (all *p* > 0.05).

## DISCUSSION

Here, we report the relationship between baseline circulating levels of eCBs/NAEs and somatosensory sensitivity to thermal and mechanical stimuli in 91 healthy participants. Thermal measures (i.e., detection thresholds, pain thresholds, or pain tolerance) and TSP were not associated with baseline eCBs/NAEs. However, PPT did show a positive relationship with circulating FAAH substrates (i.e., a composite measure comprising AEA, OEA, and PEA). Post-hoc analyses revealed that the relationship was significant for AEA and OEA, but not PEA. Furthermore, participants with either *FAAH* SNP variant (CA or AA) did not display higher circulating levels of eCBs/NAEs compared to homozygote CC carriers. Additionally, there was no association found between cannabis use, eCBs and QST measures.

Extensive preclinical work shows that cannabinoid receptors, their lipid ligands and metabolic enzymes lie at every relay site of the nociceptive neuraxis and the descending modulatory circuits [13,79,80]. Although the effects of FAAH inhibition are well documented in rodent chronic pain models, such effects have yet to be fully understood in acute pain. One study demonstrated that FAAH inhibition reversed mechanical hypersensitivity in a mouse model of acute thermal injury [81]. Another study demonstrated that pretreatment with a peripherally restricted FAAH inhibitor prevented the development of acute mechanical and thermal hypersensitivity in the carrageenan model of acute inflammation [90]. This is particularly interesting in the context of our study, given that we are measuring peripheral levels of eCBs/NAEs and their relationship with acute pain. As such, we measured baseline serum eCB/NAE levels that were not pharmacologically enhanced.

Translating preclinical findings to humans has lagged, and existing studies are fragmented across small samples, heterogeneous pain assays, and various experimental pain models. For example, one study found no association between plasma eCB levels and PPT in either patients with chronic musculoskeletal pain or healthy controls; notably, the control group in this study included only 11 participants [95]. Similarly, there was no relationship observed between circulating eCB/NAE levels and PPT in females with fibromyalgia [98] or in patients with neuromyelitis [84]. However, in another study, PPT in females with chronic widespread pain and females with chronic neck and shoulder pain was negatively correlated with microdialysate concentrations of PEA from the trapezius muscle at baseline [43].

Of note, the relationship between PPT and eCB/NAE in healthy controls was not investigated in these previous studies. Here, the association between higher FAAH-substrate levels and elevated PPT may reflect the anti-inflammatory or analgesic properties of endogenous AEA and other NAEs in deep tissues. During trapezius PPT testing, the algometer compresses the myofascial unit, activating high-threshold afferents in muscle tissue that contain nociceptive free nerve endings [70], potentially activating Piezo receptors [73]. Indeed, it is possible that FAAH substrates could reduce PPT-induced activation and transmission in nociceptive pathways: AEA and OEA are direct agonists for TRPV1 [64], whereas PEA is not [2]. Recent studies have shown that TRPV1 activation can inhibit Piezo2 activity when they are colocalized in models of visceral mechanical nociception [12,104]: thus this provides a potential downstream mechanism for AEA and OEA association with PPT, but not PEA in our sample. Clinical evidence supports this mechanism: a meta-analysis of randomized controlled trials found that exogenous PEA confers modest analgesic relief across chronic-pain conditions [4]. As such, our data suggest that individual differences in baseline AEA and OEA are associated with PPT and thus may contribute to individual differences. In contrast to the above-mentioned studies, we did not observe a relationship between PEA and PPT; the previous studies investigated changes in PEA, whereas ours focused on baseline PEA. Given the dynamic nature of eCBs/NAEs—i.e. that they are produced on demand [45,50,57,77,102], it is feasible that baseline measures of PEA are not as sensitive to individual differences in pain in healthy volunteers. Further studies are warranted, given the dearth of studies that have investigated eCBs and pain sensitivity in healthy participants. Together with the limited clinical data available, our study shows that baseline circulating levels of eCBs/NAEs are not associated with the outcome measures of standard QST tests that rely on point estimates in healthy individuals.

Rodent studies reveal sex differences in cannabinoid antinociception, with females generally showing enhanced sensitivity to cannabinoid-induced analgesia in various pain tests [26]. However, evidence regarding eCB tone is mixed, with some studies reporting that female rodents have higher AEA levels in discrete CNS regions [62], while others find higher levels in other brain regions in males [15]. In our study, females had lower WDT and HPTol, which has previously been reported in the literature [5,38,61]. However, we did not measure sex hormones, thereby not capturing previously reported cycle-dependent eCB fluctuations or progesterone– AEA interactions [19]. Interestingly, there was no association observed between circulating FAAH substrates and HPTol. This is surprising, given that AEA is also an agonist of TRPV1, the channel responsible for heat pain transduction and is activated by noxious heat (>43°C) [18]. However, baseline circulating levels of eCBs/NAEs may not be associated with point estimates of somatosensory and pain sensitivity in healthy individuals.

Contrary to expectations and what has previously been reported in the literature [69,91], carriers of at least one *FAAH* C385A allele did not have higher circulating levels of AEA, PEA or OEA, and did not differ from CC homozygotes on any thermal or mechanical sensitivity tests. However, other studies with similar representation of the *FAAH* variant in their study population have also reported no statistically significant increase in circulating eCBs/NAEs [11]. In a study of 1,000 women undergoing surgery for breast cancer, individuals with the AA genotype for the C385A *FAAH* SNP were less sensitive to cold pain pre-operatively and reported less post-operative pain than individuals with CC and AC genotypes [17]. As our sample comprised only five participants with a *FAAH* SNP AA genotype, we could not test the contribution of this genotype on QST measures with any statistical confidence. Furthermore, as eCBs/NAEs are produced on demand, *FAAH* polymorphisms may influence pain primarily when the system is challenged by stress or injury, and this may be more pronounced in AA homozygotes; thus, larger samples enriched for AA homozygotes are warranted.

Circulating eCBs are produced on-demand in response to stress [28,31,53,75], exercise [16,53], food presentation [42], inflammation and tissue injury [35,36,82], and time of day [47]. Circadian rhythms of circulating eCBs are complex and ligand-specific [47]: AEA shows a biphasic pattern peaking around 02:00 and 15:00 hours, with a nadir at 10:00 hours, while 2-AG follows a monophasic rhythm, peaking at 12:00–15:00 hours and dipping near 04:00 hours[47]. By sampling after a 12-hour fast and restricting testing to 09:00–12:00 hours, we likely captured trough values for AEA and rising, but relatively low levels for 2-AG.

Cannabis-use can also affect circulating levels of eCBs/NAEs in rodents [1]. In humans, evidence of differences in endocannabinoids in chronic users compared to non-users is mixed depending on the ligand, and the biological matrix sampled (e.g., cerebrospinal fluid, plasma, or serum) [11,56,63,71,72]. Studies investigating acute administration of exogenous cannabinoids in non-cannabis users have reported an increase in AEA and 2-AG within the following three hours of administration [97,100]. In our study, self-reported cannabis use was not associated with eCB levels or QST measures. However, we could not determine whether the amount of cannabis use was associated with differences in pain sensitivity, as our sample of those who consumed cannabis in the last 3 months was only 25 participants. Future studies should directly measure phytocannabinoids and their metabolites in larger samples, across biological matrices to better determine the impact of cannabis usage on eCB/NAE levels and pain sensitivity.

Several methodological limitations warrant consideration. Our cross-sectional design precludes causal inference regarding eCB-QST relationships. It is possible that a more comprehensive analysis that focuses solely on thermal pain sensitivity with multiple suprathreshsold stimuli at multiple sites might reveal an associated with eCBs/NAEs. Also, the unequal distribution of the *FAAH* genotypes in our sample limits our power to detect the effects of this genotype on QST responses. Additionally, the correlation between circulating serum concentrations and synaptic eCB levels within spinal or supraspinal pain circuits remains unknown [49]. Current non-invasive methodologies cannot adequately capture central nervous system eCB dynamics, as spinal tap to sample cerebrospinal fluid is associated with risks that are unjustifiable in healthy people. Furthermore, extant ligands for pharmacological brain imaging (i.e., positron-emission tomography), such as those developed for FAAH and CB^1^ receptor, bind irreversibly [101]. Lastly, cannabis exposure assessment relied on self-report rather than toxicological confirmation, introducing potential reporting bias, and limited resolution.

Future studies are required to determine: (i) whether dynamic biochemistry is more sensitive to eCB/NAE-pain relationships—coupling serial blood or interstitial-fluid sampling to QST and intervention-based tasks could be more sensitive to eCB/NAE-pain relationships; and (ii) causal relationships between eCBs/NAEs and pain using pharmacological challenges (e.g. with FAAH inhibitors or CB^1^ receptor antagonists).

Taken together, our data indicate that baseline eCBs and FAAH genotype are not associated with the outcome measures of standard QST tests that rely on point estimates in healthy adults; nonetheless, higher circulating FAAH substrates are associated with higher PPT. These findings refine our mechanistic understanding of the human eCB system and its relationship to acute pain sensitivity and highlight the importance of pain-modality test, dynamic ligand sampling, and individual biochemical profiling when designing eCB-based analgesic strategies.

## ACKNOWLEDGEMENTS

The authors have no conflicts of interest to declare. This work was supported by a Canadian Institutes of Health Research (CIHR) Project Grant awarded to M. M., D. P. F. and L. Y. A. (183703). M. M. holds a Canada Research Chair (Tier 2) in Pain NeuroImaging, and is supported by a University of Toronto Pain Scientist Award. L. Y. A. was supported in part by the Intramural Research Program of the National Institutes of Health (NIH; ZIA-AT000030). The contributions of the NIH author were made as part of their official duties as NIH federal employees, are in compliance with agency policy requirements, and are considered Works of the United States Government. However, the findings and conclusions presented in this paper are those of the author(s) and do not necessarily reflect the views of the NIH or the U.S. Department of Health and Human Services. S. B. was funded by a Government of Ireland Postgraduate Research Scholarship from the Irish Research Council (GOIPG/2019/3945). S. S. A. was supported by an Ontario Graduate Scholarship (OGS), a Natural Sciences and Engineering Research Council (NSERC) Canada Graduate Scholarship – Masters (CGS-M), and the University of Toronto’s Faculty of Dentistry Harron Fund Award. M. A. C. was supported by an NSERC CGS-M. R. T. was supported by University of Toronto’s Faculty of Dentistry Harron Fund Award, and an OGS.

Dr. Liat Honigman, who is a co-author on this paper, was a beloved lab member and research associate in the Centre for Multimodal Sensorimotor and Pain Research (CMSPR) at the University of Toronto. She died on March 28, 2024. This, and many other projects in the lab would not have been possible without her expertise, guidance, and support. Liat was kind, smart, and a mentor to many trainees in the CMSPR. She held the highest standards for science, but believed in a compassionate approach to research. She is truly missed.

Data Accessibility: All data for this manuscript is appended in Supplementary Materials.

## Author Contributions

Conception of the study: MM, LYA and DPF

Design of the Study: LH, IB and MM, DPF

Data Acquisition: SAF, SB, NM, CS, MAC, RT

Data Analysis: SSA, SAF, SB, KM

Interpretation of the data: SAF, SSA, MM, LYA, DPF

Drafting of manuscript: SAF, SSA, MM

Reviewing and Editing of Manuscript and Final Approval: SAF, SSA, SB, KM, NM, RT, CS, MAC, IB, LYA, DPF, MM

