## Supplementary material for "The Contribution of Baseline Circulating Endocannabinoids to Individual Differences in Human Pain Sensitivity: A Quantitative Sensory Testing Study": Figure S

^1^Centre for Multimodal Sensorimotor and Pain Research, Faculty of Dentistry, University of Toronto, Toronto, Canada;^2^Pharmacology and Therapeutics, and ^3^Centre for Pain Research, University of Galway, Galway, Ireland; ^4^Centre for Addiction and Mental Health, Toronto, ON, Canada; ^5^National Center For Complementary and Integrative Health, ^6^National Institute of Drug Abuse, and ^7^National Institutes of Mental Health, National Institutes of Health, Bethesda, MD, United States; ^8^University of Toronto Centre for the Study of Pain, Toronto, ON, Canada;  ^9^Krembil Research Institute, University Health Network, Toronto, ON, Canada

*^Co-First Authors; ^†^Co-Senior Authors*

Table S1: Frequency and Count of *FAAH* C385A (rs324420) alleles

| Geographic Region | Genotype | | | | | |
| --- | --- | --- | --- | --- | --- | --- |
|  | CC | | AC | | AA | |
|  | Frequency | Count | Frequency | Count | Frequency | Count |
| All | 0.555 | 1390 | 0.367 | 918 | 0.078 | 196 |
| African | 0.404 | 267 | 0.457 | 302 | 0.139 | 92 |
| American | 0.427 | 148 | 0.444 | 154 | 0.13 | 45 |
| East Asian | 0.679 | 342 | 0.292 | 147 | 0.03 | 15 |
| European | 0.622 | 313 | 0.334 | 168 | 0.044 | 22 |
| South Asian | 0.654 | 320 | 0.301 | 147 | 0.045 | 22 |
| Source: Ensembl database [1] | | | | | | |

Table S2. Comparison of eCB/NAE levels in males and females

| eCB/NAE Ligand | Shapiro-Wilk’s Test | | Independent Samples Group Comparison | | Mean ± SD | | Effect Size | |
| --- | --- | --- | --- | --- | --- | --- | --- | --- |
|  | *W* | *p* | T/U* | *p* | *Males* | *Females* | | *g*/*rb*** |
| AEA | 0.91 | 6.17E-06 | 1053 | 0.76 | 2.27 ± 0.79 | 2.32 ± 0.79 | | 0.04 |
| 2-AG | 0.75 | 4.55E-11 | 932 | 0.51 | 3.91 ± 1.91 | 4.32 ± 3.43 | | -0.08 |
| OEA | 0.92 | 2.54E-5 | 1206 | 0.13 | 25.07 ± 8.21 | 28.30 ± 10.11 | | 0.19 |
| PEA | 0.99 | 0.43 | -1.036 | 0.30 | 13.88 ± 3.48 | 13.08 ± 3.83 | | -0.22 |

*Student’s t-tests performed for normally distributed variables, and Mann-Whitney *U-*tests performed for non-normally distributed variables

**Effect size is given by Hedge’s g for Student’s t-test and rank biserial correlation for Mann-Whitney test.

Table S3. Effects of *FAAH* SNP on eCB/NAE levels

| eCB/NAE Ligand | Shapiro-Wilk’s Test | | Independent Samples Group Comparison* | | Mean ± SD | | Effect Size |
| --- | --- | --- | --- | --- | --- | --- | --- |
|  | *W* | *p* | T/U** | *p* | *CC* | *A–* | *g*/*rb**** |
| AEA | 0.90 | 5.71E-6 | 883 | 0.88 | 2.29 ± 0.82 | 2.31 ± 0.74 | -0.07 |
| 2-AG | 0.74 | 2.16E-11 | 883 | 0.88 | 4.17 ± 3.21 | 4.07 ± 2.16 | 0.07 |
| OEA | 0.92 | 1.87E-5 | 737 | 0.09 | 12.91 ± 3.44 | 14.36 ± 3.98 | -0.22 |
| PEA | 0.99 | 0.44 | -1.81 | 0.07 | 25.65 ± 8.69 | 29.23 ± 10.4 | -0.40 |

*Grouping variable is a binary: presence/absence of A allele

**Student’s t-tests performed for normally distributed variables, and Mann-Whitney *U-*tests performed for non-normally distributed variables

***Effect size is given by Hedge’s g for Student’s t-test and rank biserial correlation for Mann-Whitney test.

Table S4. Factor Reduction for FAAH Substrate Component Loading Table

|  | Component Loadings | Uniqueness |
| --- | --- | --- |
| OEA | 0.91 | 0.177 |
| AEA | 0.88 | 0.230 |
| PEA | 0.77 | 0.404 |

Table S5a. Regression Model ANOVA Table for WDT

| Model | | AIC |  | | Sum of Squares | | df | | Mean Square | | F | | *p* |
| --- | --- | --- | --- | --- | --- | --- | --- | --- | --- | --- | --- | --- | --- |
| M₀ |  | 249.05 | Regression |  | 3.782 |  | 2 |  | 1.891 |  | 2.080 |  | 0.131 |
|  |  |  | Residual |  | 78.194 |  | 86 |  | 0.909 |  |  |  |  |
|  |  |  | Total |  | 81.976 |  | 88 |  |  |  |  |  |  |
| M₁ |  | 243.04 | Regression |  | 10.513 |  | 3 |  | 3.504 |  | 4.168 |  | 0.008 |
|  |  |  | Residual |  | 71.463 |  | 85 |  | 0.841 |  |  |  |  |
|  |  |  | Total |  | 81.976 |  | 88 |  |  |  |  |  |  |
| M₂ |  | 245.03 | Regression |  | 10.519 |  | 4 |  | 2.630 |  | 3.091 |  | 0.020 |
|  |  |  | Residual |  | 71.458 |  | 84 |  | 0.851 |  |  |  |  |
|  |  |  | Total |  | 81.976 |  | 88 |  |  |  |  |  |  |
| M₃ |  | 248.78 | Regression |  | 10.724 |  | 6 |  | 1.787 |  | 2.057 |  | 0.067 |
|  |  |  | Residual |  | 71.252 |  | 82 |  | 0.869 |  |  |  |  |
|  |  |  | Total |  | 81.976 |  | 88 |  |  |  |  |  |  |
| M₄ |  | 252.49 | Regression |  | 10.954 |  | 8 |  | 1.369 |  | 1.542 |  | 0.156 |
|  |  |  | Residual |  | 71.022 |  | 80 |  | 0.888 |  |  |  |  |
|  |  |  | Total |  | 81.976 |  | 88 |  |  |  |  |  |  |
| Note.  M₀ includes Age, BMI | | | | | | | | | | | | | |
| Note.  M₁ includes Age, BMI, Sex | | | | | | | | | | | | | |
| Note.  M₂ includes Age, BMI, Sex, FAAH SNP Genotype | | | | | | | | | | | | | |
| Note.  M₃ includes Age, BMI, Sex, FAAH SNP Genotype, 2-AG, FAAH Substrates | | | | | | | | | | | | | |
| Note.  M₄ includes Age, BMI, 2-AG, FAAH Substrates, Sex, FAAH SNP Genotype, Cannabis Usage | | | | | | | | | | | | | |

Table S5b. Coefficients Table of Winning Model for WDT

| Predictor | Unstandardized | Standard Error | Standardized^a^ | *t* | *p* |
| --- | --- | --- | --- | --- | --- |
| (Intercept) | -0.008 | 0.872 |  | -0.009 | 0.993 |
| Age | 0.042 | 0.021 | 0.211 | 1.977 | 0.051 |
| BMI | -0.031 | 0.035 | -0.097 | -0.884 | 0.379 |
| Sex | -0.578 | 0.204 |  | -2.830 | 0.006 |
| ^a^Standardized coefficients can only be computed for continuous predictors. | | | | | |

Table S6. Regression Model ANOVA Table for CDT

| Model | | AIC |  | | Sum of Squares | | df | | Mean Square | | F | | *p* |
| --- | --- | --- | --- | --- | --- | --- | --- | --- | --- | --- | --- | --- | --- |
| M₀ |  | 250.82 | Regression |  | 4.685 |  | 2 |  | 2.342 |  | 2.526 |  | 0.086 |
|  |  |  | Residual |  | 79.761 |  | 86 |  | 0.927 |  |  |  |  |
|  |  |  | Total |  | 84.445 |  | 88 |  |  |  |  |  |  |
| M₁ |  | 252.76 | Regression |  | 4.731 |  | 3 |  | 1.577 |  | 1.681 |  | 0.177 |
|  |  |  | Residual |  | 79.715 |  | 85 |  | 0.938 |  |  |  |  |
|  |  |  | Total |  | 84.445 |  | 88 |  |  |  |  |  |  |
| M₂ |  | 249.98 | Regression |  | 8.902 |  | 4 |  | 2.226 |  | 2.475 |  | 0.050 |
|  |  |  | Residual |  | 75.543 |  | 84 |  | 0.899 |  |  |  |  |
|  |  |  | Total |  | 84.445 |  | 88 |  |  |  |  |  |  |
| M₃ |  | 254.42 | Regression |  | 9.962 |  | 6 |  | 1.660 |  | 1.828 |  | 0.104 |
|  |  |  | Residual |  | 74.484 |  | 82 |  | 0.908 |  |  |  |  |
|  |  |  | Total |  | 84.445 |  | 88 |  |  |  |  |  |  |
| M₄ |  | 258.04 | Regression |  | 10.257 |  | 8 |  | 1.282 |  | 1.383 |  | 0.217 |
|  |  |  | Residual |  | 74.188 |  | 80 |  | 0.927 |  |  |  |  |
|  |  |  | Total |  | 84.445 |  | 88 |  |  |  |  |  |  |
| Note.  M₀ includes Age, BMI | | | | | | | | | | | | | |
| Note.  M₁ includes Age, BMI, Sex | | | | | | | | | | | | | |
| Note.  M₂ includes Age, BMI, Sex, FAAH SNP Genotype | | | | | | | | | | | | | |
| Note.  M₃ includes Age, BMI, Sex, FAAH SNP Genotype, 2-AG, FAAH Substrates | | | | | | | | | | | | | |
| Note.  M₄ includes Age, BMI, 2-AG, FAAH Substrates, Sex, FAAH SNP Genotype, Cannabis Usage | | | | | | | | | | | | | |

Table S7. Regression Model ANOVA Table for HPT

|  | |  | | | | | | | | | | | |
| --- | --- | --- | --- | --- | --- | --- | --- | --- | --- | --- | --- | --- | --- |
| Model | | AIC |  | | Sum of Squares | | df | | Mean Square | | F | | *p* |
| M₀ |  | 262.10 | Regression |  | 5.801 |  | 2 |  | 2.900 |  | 2.935 |  | 0.058 |
|  |  |  | Residual |  | 86.949 |  | 88 |  | 0.988 |  |  |  |  |
|  |  |  | Total |  | 92.749 |  | 90 |  |  |  |  |  |  |
| M₁ |  | 264.05 | Regression |  | 5.854 |  | 3 |  | 1.951 |  | 1.954 |  | 0.127 |
|  |  |  | Residual |  | 86.896 |  | 87 |  | 0.999 |  |  |  |  |
|  |  |  | Total |  | 92.749 |  | 90 |  |  |  |  |  |  |
| M₂ |  | 264.68 | Regression |  | 7.145 |  | 4 |  | 1.786 |  | 1.794 |  | 0.137 |
|  |  |  | Residual |  | 85.605 |  | 86 |  | 0.995 |  |  |  |  |
|  |  |  | Total |  | 92.749 |  | 90 |  |  |  |  |  |  |
| M₃ |  | 267.06 | Regression |  | 8.657 |  | 6 |  | 1.443 |  | 1.441 |  | 0.209 |
|  |  |  | Residual |  | 84.092 |  | 84 |  | 1.001 |  |  |  |  |
|  |  |  | Total |  | 92.749 |  | 90 |  |  |  |  |  |  |
| M₄ |  | 266.82 | Regression |  | 12.486 |  | 8 |  | 1.561 |  | 1.595 |  | 0.139 |
|  |  |  | Residual |  | 80.263 |  | 82 |  | 0.979 |  |  |  |  |
|  |  |  | Total |  | 92.749 |  | 90 |  |  |  |  |  |  |
| Note.  M₀ includes Age, BMI | | | | | | | | | | | | | |
| Note.  M₁ includes Age, BMI, Sex | | | | | | | | | | | | | |
| Note.  M₂ includes Age, BMI, Sex, FAAH SNP Genotype | | | | | | | | | | | | | |
| Note.  M₃ includes Age, BMI, Sex, FAAH SNP Genotype, 2-AG, FAAH Substrates | | | | | | | | | | | | | |
| Note.  M₄ includes Age, BMI, 2-AG, FAAH Substrates, Sex, FAAH SNP Genotype, Cannabis Usage | | | | | | | | | | | | | |

| Table S8. Regression Model ANOVA Table for CPT | | | | | | | | | | | | | |
| --- | --- | --- | --- | --- | --- | --- | --- | --- | --- | --- | --- | --- | --- |
| Model | | AIC |  | | Sum of Squares | | df | | Mean Square | | F | | p |
| M₀ |  | 263.92 | Regression |  | 0.930 |  | 2 |  | 0.465 |  | 0.461 |  | 0.632 |
|  |  |  | Residual |  | 88.701 |  | 88 |  | 1.008 |  |  |  |  |
|  |  |  | Total |  | 89.632 |  | 90 |  |  |  |  |  |  |
| M₁ |  | 265.85 | Regression |  | 0.995 |  | 3 |  | 0.332 |  | 0.326 |  | 0.807 |
|  |  |  | Residual |  | 88.636 |  | 87 |  | 1.019 |  |  |  |  |
|  |  |  | Total |  | 89.632 |  | 90 |  |  |  |  |  |  |
| M₂ |  | 267.85 | Regression |  | 0.999 |  | 4 |  | 0.250 |  | 0.242 |  | 0.914 |
|  |  |  | Residual |  | 88.633 |  | 86 |  | 1.031 |  |  |  |  |
|  |  |  | Total |  | 89.632 |  | 90 |  |  |  |  |  |  |
| M₃ |  | 271.16 | Regression |  | 1.670 |  | 6 |  | 0.278 |  | 0.266 |  | 0.951 |
|  |  |  | Residual |  | 87.962 |  | 84 |  | 1.047 |  |  |  |  |
|  |  |  | Total |  | 89.632 |  | 90 |  |  |  |  |  |  |
| M₄ |  | 268.47 | Regression |  | 7.901 |  | 8 |  | 0.988 |  | 0.991 |  | 0.449 |
|  |  |  | Residual |  | 81.731 |  | 82 |  | 0.997 |  |  |  |  |
|  |  |  | Total |  | 89.632 |  | 90 |  |  |  |  |  |  |
| Note.  M₀ includes Age, BMI | | | | | | | | | | | | | |
| Note.  M₁ includes Age, BMI, Sex | | | | | | | | | | | | | |
| Note.  M₂ includes Age, BMI, Sex, FAAH SNP Genotype | | | | | | | | | | | | | |
| Note.  M₃ includes Age, BMI, 2-AG, FAAH Substrates, Sex, FAAH SNP Genotype | | | | | | | | | | | | | |
| Note.  M₄ includes Age, BMI, 2-AG, FAAH Substrates, Sex, FAAH SNP Genotype, Cannabis Usage | | | | | | | | | | | | | |

Table S9a. Regression Model ANOVA Table for HPTol

| Model | | AIC |  | | Sum of Squares | | df | | Mean Square | | F | | p |
| --- | --- | --- | --- | --- | --- | --- | --- | --- | --- | --- | --- | --- | --- |
| M₀ |  | 234.36 | Regression |  | 4.701 |  | 2 |  | 2.350 |  | 2.363 |  | 0.101 |
|  |  |  | Residual |  | 77.571 |  | 78 |  | 0.995 |  |  |  |  |
|  |  |  | Total |  | 82.272 |  | 80 |  |  |  |  |  |  |
| M₁ |  | 227.60 | Regression |  | 12.652 |  | 3 |  | 4.217 |  | 4.664 |  | 0.005 |
|  |  |  | Residual |  | 69.620 |  | 77 |  | 0.904 |  |  |  |  |
|  |  |  | Total |  | 82.272 |  | 80 |  |  |  |  |  |  |
| M₂ |  | 227.48 | Regression |  | 14.457 |  | 4 |  | 3.614 |  | 4.050 |  | 0.005 |
|  |  |  | Residual |  | 67.815 |  | 76 |  | 0.892 |  |  |  |  |
|  |  |  | Total |  | 82.272 |  | 80 |  |  |  |  |  |  |
| M₃ |  | 231.35 | Regression |  | 14.563 |  | 6 |  | 2.427 |  | 2.653 |  | 0.022 |
|  |  |  | Residual |  | 67.709 |  | 74 |  | 0.915 |  |  |  |  |
|  |  |  | Total |  | 82.272 |  | 80 |  |  |  |  |  |  |
| M₄ |  | 232.57 | Regression |  | 16.850 |  | 8 |  | 2.106 |  | 2.318 |  | 0.028 |
|  |  |  | Residual |  | 65.422 |  | 72 |  | 0.909 |  |  |  |  |
|  |  |  | Total |  | 82.272 |  | 80 |  |  |  |  |  |  |
| Note.  M₀ includes Age, BMI | | | | | | | | | | | | | |
| Note.  M₁ includes Age, BMI, Sex | | | | | | | | | | | | | |
| Note.  M₂ includes Age, BMI, Sex, FAAH SNP Genotype | | | | | | | | | | | | | |
| Note.  M₃ includes Age, BMI, Sex, FAAH SNP Genotype, 2-AG, FAAH Substrates | | | | | | | | | | | | | |
| Note.  M₄ includes Age, BMI, 2-AG, FAAH Substrates, Sex, FAAH SNP Genotype, Cannabis Usage | | | | | | | | | | | | | |

Table S9b. Regression Model Coefficients Table for HPTol

|  | | Unstandardized | | Standard Error | | Standardizedᵃ | | t | | p |
| --- | --- | --- | --- | --- | --- | --- | --- | --- | --- | --- |
| (Intercept) |  | -0.852 |  | 0.940 |  |  |  | -0.907 |  | 0.367 |
| Age |  | 0.007 |  | 0.023 |  | 0.033 |  | 0.295 |  | 0.769 |
| BMI |  | 0.040 |  | 0.039 |  | 0.120 |  | 1.037 |  | 0.303 |
| Sex (Female) |  | -0.635 |  | 0.221 |  |  |  | -2.874 |  | 0.005 |
| FAAH SNP genotype (1) |  | 0.325 |  | 0.229 |  |  |  | 1.422 |  | 0.159 |
| ^a^Standardized coefficients can only be computed for continuous predictors. | | | | | | | | | | |

Table S10. Regression Model ANOVA Table for CPTol

| Model | | AIC |  | | Sum of Squares | | df | | Mean Square | | F | | p |
| --- | --- | --- | --- | --- | --- | --- | --- | --- | --- | --- | --- | --- | --- |
| M₀ |  | 1002.53 | Regression |  | 9165.240 |  | 2 |  | 4582.620 |  | 1.357 |  | 0.263 |
|  |  |  | Residual |  | 297105.981 |  | 88 |  | 3376.204 |  |  |  |  |
|  |  |  | Total |  | 306271.221 |  | 90 |  |  |  |  |  |  |
| M₁ |  | 1003.43 | Regression |  | 12727.214 |  | 3 |  | 4242.405 |  | 1.257 |  | 0.294 |
|  |  |  | Residual |  | 293544.006 |  | 87 |  | 3374.069 |  |  |  |  |
|  |  |  | Total |  | 306271.221 |  | 90 |  |  |  |  |  |  |
| M₂ |  | 1005.40 | Regression |  | 12817.733 |  | 4 |  | 3204.433 |  | 0.939 |  | 0.445 |
|  |  |  | Residual |  | 293453.488 |  | 86 |  | 3412.250 |  |  |  |  |
|  |  |  | Total |  | 306271.221 |  | 90 |  |  |  |  |  |  |
| M₃ |  | 1006.46 | Regression |  | 22149.433 |  | 6 |  | 3691.572 |  | 1.091 |  | 0.374 |
|  |  |  | Residual |  | 284121.788 |  | 84 |  | 3382.402 |  |  |  |  |
|  |  |  | Total |  | 306271.221 |  | 90 |  |  |  |  |  |  |
| M₄ |  | 1008.75 | Regression |  | 27429.101 |  | 8 |  | 3428.638 |  | 1.008 |  | 0.436 |
|  |  |  | Residual |  | 278842.119 |  | 82 |  | 3400.514 |  |  |  |  |
|  |  |  | Total |  | 306271.221 |  | 90 |  |  |  |  |  |  |
| Note.  M₀ includes Age, BMI | | | | | | | | | | | | | |
| Note.  M₁ includes Age, BMI, Sex | | | | | | | | | | | | | |
| Note.  M₂ includes Age, BMI, Sex, FAAH SNP Genotype | | | | | | | | | | | | | |
| Note.  M₃ includes Age, BMI, Sex, FAAH SNP Genotype, 2-AG, FAAH Substrates | | | | | | | | | | | | | |
| Note.  M₄ includes Age, BMI, 2-AG, FAAH Substrates, Sex, FAAH SNP Genotype, Cannabis Usage | | | | | | | | | | | | | |

Table S11a. Regression Model ANOVA Table for PPT

|  | |  | | | | | | | | | | | |
| --- | --- | --- | --- | --- | --- | --- | --- | --- | --- | --- | --- | --- | --- |
| Model | | AIC |  | | Sum of Squares | | df | | Mean Square | | F | | p |
| M₀ |  | 1205.56 | Regression |  | 318651.807 |  | 2 |  | 159325.904 |  | 5.069 |  | 0.008 |
|  |  |  | Residual |  | 2.766×10^+6^ |  | 88 |  | 31433.175 |  |  |  |  |
|  |  |  | Total |  | 3.085×10^+6^ |  | 90 |  |  |  |  |  |  |
| M₁ |  | 1206.56 | Regression |  | 348929.714 |  | 3 |  | 116309.905 |  | 3.699 |  | 0.015 |
|  |  |  | Residual |  | 2.736×10^+6^ |  | 87 |  | 31446.454 |  |  |  |  |
|  |  |  | Total |  | 3.085×10^+6^ |  | 90 |  |  |  |  |  |  |
| M₂ |  | 1208.07 | Regression |  | 363595.383 |  | 4 |  | 90898.846 |  | 2.873 |  | 0.028 |
|  |  |  | Residual |  | 2.721×10^+6^ |  | 86 |  | 31641.579 |  |  |  |  |
|  |  |  | Total |  | 3.085×10^+6^ |  | 90 |  |  |  |  |  |  |
| M₃ |  | 1202.65 | Regression |  | 631240.596 |  | 6 |  | 105206.766 |  | 3.602 |  | 0.003 |
|  |  |  | Residual |  | 2.454×10^+6^ |  | 84 |  | 29208.698 |  |  |  |  |
|  |  |  | Total |  | 3.085×10^+6^ |  | 90 |  |  |  |  |  |  |
| M₄ |  | 1206.20 | Regression |  | 643258.146 |  | 8 |  | 80407.268 |  | 2.701 |  | 0.011 |
|  |  |  | Residual |  | 2.442×10^+6^ |  | 82 |  | 29774.549 |  |  |  |  |
|  |  |  | Total |  | 3.085×10^+6^ |  | 90 |  |  |  |  |  |  |
| Note.  M₀ includes Age, BMI | | | | | | | | | | | | | |
| Note.  M₁ includes Age, BMI, Sex | | | | | | | | | | | | | |
| Note.  M₂ includes Age, BMI, Sex, FAAH SNP Genotype | | | | | | | | | | | | | |
| Note.  M₃ includes Age, BMI, Sex, FAAH SNP Genotype, FAAH Substrates, 2-AG | | | | | | | | | | | | | |
| Note.  M₄ includes Age, BMI, FAAH Substrates, 2-AG, Sex, FAAH SNP Genotype, Cannabis Usage | | | | | | | | | | | | | |

Table S11b. Regression Model Coefficients Table for PPT

|  | Unstandardized | Standard Error | Standardized | t | p |
| --- | --- | --- | --- | --- | --- |
| (Intercept) | 72.35 | 165.56 |  | 0.437 | 0.663 |
| Age | 6.11 | 4.05 | 0.160 | 1.508 | 0.135 |
| BMI | 9.59 | 6.66 | 0.156 | 1.441 | 0.153 |
| Sex | -43.27 | 37.82 |  | -1.144 | 0.256 |
| FAAH SNP genotype | 10.68 | 38.84 |  | 0.275 | 0.784 |
| FAAH Substrates | 57.95 | 19.54 | 0.314 | 2.966 | 0.004 |
| 2-AG | -9.84 | 6.81 | -0.153 | -1.445 | 0.152 |
| ^a^ Standardized coefficients can only be computed for continuous predictors. | | | | | |

Table S12. Regression Model ANOVA Table for TSP

| Model | | AIC |  | | Sum of Squares | | df | | Mean Square | | F | | p |
| --- | --- | --- | --- | --- | --- | --- | --- | --- | --- | --- | --- | --- | --- |
| M₀ |  | 720.72 | Regression |  | 392.347 |  | 2 |  | 196.174 |  | 1.286 |  | 0.282 |
|  |  |  | Residual |  | 13427.464 |  | 88 |  | 152.585 |  |  |  |  |
|  |  |  | Total |  | 13819.811 |  | 90 |  |  |  |  |  |  |
| M₁ |  | 720.91 | Regression |  | 656.663 |  | 3 |  | 218.888 |  | 1.447 |  | 0.235 |
|  |  |  | Residual |  | 13163.148 |  | 87 |  | 151.301 |  |  |  |  |
|  |  |  | Total |  | 13819.811 |  | 90 |  |  |  |  |  |  |
| M₂ |  | 720.36 | Regression |  | 1019.893 |  | 4 |  | 254.973 |  | 1.713 |  | 0.154 |
|  |  |  | Residual |  | 12799.919 |  | 86 |  | 148.836 |  |  |  |  |
|  |  |  | Total |  | 13819.811 |  | 90 |  |  |  |  |  |  |
| M₃ |  | 721.33 | Regression |  | 1439.850 |  | 6 |  | 239.975 |  | 1.628 |  | 0.149 |
|  |  |  | Residual |  | 12379.962 |  | 84 |  | 147.380 |  |  |  |  |
|  |  |  | Total |  | 13819.811 |  | 90 |  |  |  |  |  |  |
| M₄ |  | 723.57 | Regression |  | 1676.572 |  | 8 |  | 209.572 |  | 1.415 |  | 0.203 |
|  |  |  | Residual |  | 12143.239 |  | 82 |  | 148.088 |  |  |  |  |
|  |  |  | Total |  | 13819.811 |  | 90 |  |  |  |  |  |  |
| Note.  M₀ includes Age, BMI | | | | | | | | | | | | | |
| Note.  M₁ includes Age, BMI, Sex | | | | | | | | | | | | | |
| Note.  M₂ includes Age, BMI, Sex, FAAH SNP Genotype | | | | | | | | | | | | | |
| Note.  M₃ includes Age, BMI, Sex, FAAH SNP Genotype, 2-AG, FAAH Substrates | | | | | | | | | | | | | |
| Note.  M₄ includes Age, BMI, 2-AG, FAAH Substrates, Sex, FAAH SNP Genotype, Cannabis Usage | | | | | | | | | | | | | |
